# Chlorophyll content and chlorophyll fluorescence as physiological parameters for monitoring *Orobanche foetida* Poir. infestation on faba bean (*Vicia faba* L.)

**DOI:** 10.1101/2020.10.19.345173

**Authors:** Moez Amri, Zouhaier Abbes, Imen Trabelsi, Michel Edmond Ghanem, Rachid Mentag, Mohamed Kharrat

## Abstract

In total, 39 faba bean (*Vicia faba* L.) advanced lines were evaluated for resistance to broomrape *Orobanche foetida* under highly infested field conditions. The trials were conducted during two consecutive copping seasons at Oued-Beja Research Station in Tunisia. The advanced lines XAR-VF00.13-1-2-1-2-1 and XBJ90.04-2-3-1-1-1-2A expressed high resistance level to *O. foetida* exceeding those recorded for resistance checks Najeh and Baraca. Results showed that *O. foetida* significantly affected the biomass, grain yield, chlorophyll content index (CCI) and the maximum quantum efficiency (F_v_/F_m_ ratio). No significant effect of *O. foetida* parasitism was observed on host plant water content (WC). *O. foetida* parasitism significantly affected both CCI and F_v_/F_m_ ratio. CCI decreases varied from 46.4% for the susceptible check Badi and 4.2% and 9.3% observed for the genotypes Baraca and XBJ90.04-2-3-1-1-1-2A. Compared to susceptible check, slight decreases of F_v_/F_m_ ratio were observed for both advanced lines XBJ90.04-2-3-1-1-1-2A and XAR-VF00.13-1-2-1-2-1. Correlation between CCI and F_v_/F_m_ with the resistance to broomrape makes this, easy-to-measure, parameter very useful as a practical screening tool for early parasitism detection, diagnosis and identification and selection of high resistant plants against this pathogen.

## Introduction

Broomrapes (*Orobanche* spp.) are holoparasitic plants completely dependent on the host for their nutritional requirements. In the Mediterranean region, where broomrapes are considered as a serious threat, *Orobanche* causes important damages and yield losses on many legume crops [1, 2, 3]. In Tunisia, *Orobanche foetida, O. crenata, O. cumana,* and *Phelipanche ramosa* were found parasitizing many crops such as faba bean, chickpea, lentil, grass pea, sunflower, [4, 5]. While *O. crenata* was mentioned as a serious pest for decades, *O. foetida* has been presented as an emerging problem for many legume crops such as faba bean, chickpea, lentil, grass pea, medick, common and narbon vetch [4, 6, 7]. The *Orobanche* infested area in Tunisia is estimated now to more than 80,000 ha mostly situated in the main grain legumes production area (data non-published). As a result, in high infested fields, farmers abandoned planting legumes especially faba bean which were substituted by non-host crops such as wheat leading to a strict wheat mono-cropping system. The detrimental effect of *Orobanche* is associated with their high seed viability (up to 15-20 years) and multiplication rate. In order to slow down and stop the fast spread of the parasite from invading new agricultural lands, control strategies and preventive actions should target decreasing of *Orobanche* seed bank in the soil and minimizing new seed production [8]. Till date, no single control method/technology has shown successful, and all control strategies resulted in an incomplete protection of the crop [9, 10, 11]. The only effective method to control *Orobanche* is through an integrated management approach that should be based, amongst others, on genetic resistance. Farmers should use resistant varieties and avoid planting contaminated seeds, spreading contaminated manure and soil, control animals grazing and avoid moving contaminated machinery from infested to non-infested fields [12].

Research is needed for generating new technologies and developing new resistant varieties and effective screening tools. Many resistance mechanisms were studied focusing mainly on the physical and biochemical host-parasite interface including *Orobanche* seed germination stimulant and inhibitors, host plant roots physical barrier and root architecture [2, 13, 14, 15]. While avoidance of dispersal of broomrape is virtually difficult, crop resistance and prevention measures could be the most effective and economical methods to reduce this root parasitic weed infestations. Genetic resistance coupled with other control methods result someway in good control of the parasite with significant decreases of the damages. Such integrated control strategy could be improved through early detection and monitoring of the underground infestation and the parasite development. Chlorophyll fluorescence, which is a non-destructive and rapid assessing mean of photochemical quantum yield and photoinhibition, could be used for early *Orobanche* infestation and estimate its impact on the host plant. It is widely used as a plant response indicator under many abiotic and biotic constraints such as heat, drought, waterlogging, salt stress, nitrogen deficiency, pathogen infection and herbicide resistance [16, 17, 18]. However, only few studies were conducted on parasitism effect on host plant chlorophyll fluorescence [19, 20, 21]. As reported by Maxwell and Johnson [22], the photochemical processes alterations are usually the first signs in the stressed plant leaves that could be used to estimate photosynthetic performance under stress conditions. These photochemical processes alterations appear in the chlorophyll fluorescence kinetics and induce changes in the established fluorescence parameters and consequently PSII damages.

The purpose of this study was to evaluate the response of 39 faba bean advanced lines to *O. foetida* parasitism, identify potential resistance sources and study the effect of the parasite on plant growth and seed yield in correlation with chlorophyll content and chlorophyll fluorescence.

## Material and Methods

### Plant material and field trials

A set of 39 small-seeded faba bean advanced lines, developed from crosses performed in Tunisia, were used for a first-year (2009/2010) screening and evaluation for resistance to *O. foetida*. Three checks were added to the list, two Tunisian varieties Badi and Najeh and a Spanish variety Baraca. Both varieties Najeh and Baraca, carrying partial resistance to *O. foetida* and *O. crenata* [23, 1] were used as resistance check while Badi was used as susceptible check (Table 1). The screening was performed under high *O. foetida* infested sick plot at Oued-Beja Research Station - Tunisia (36°44’N; 9°13’E).

**Table 1:**
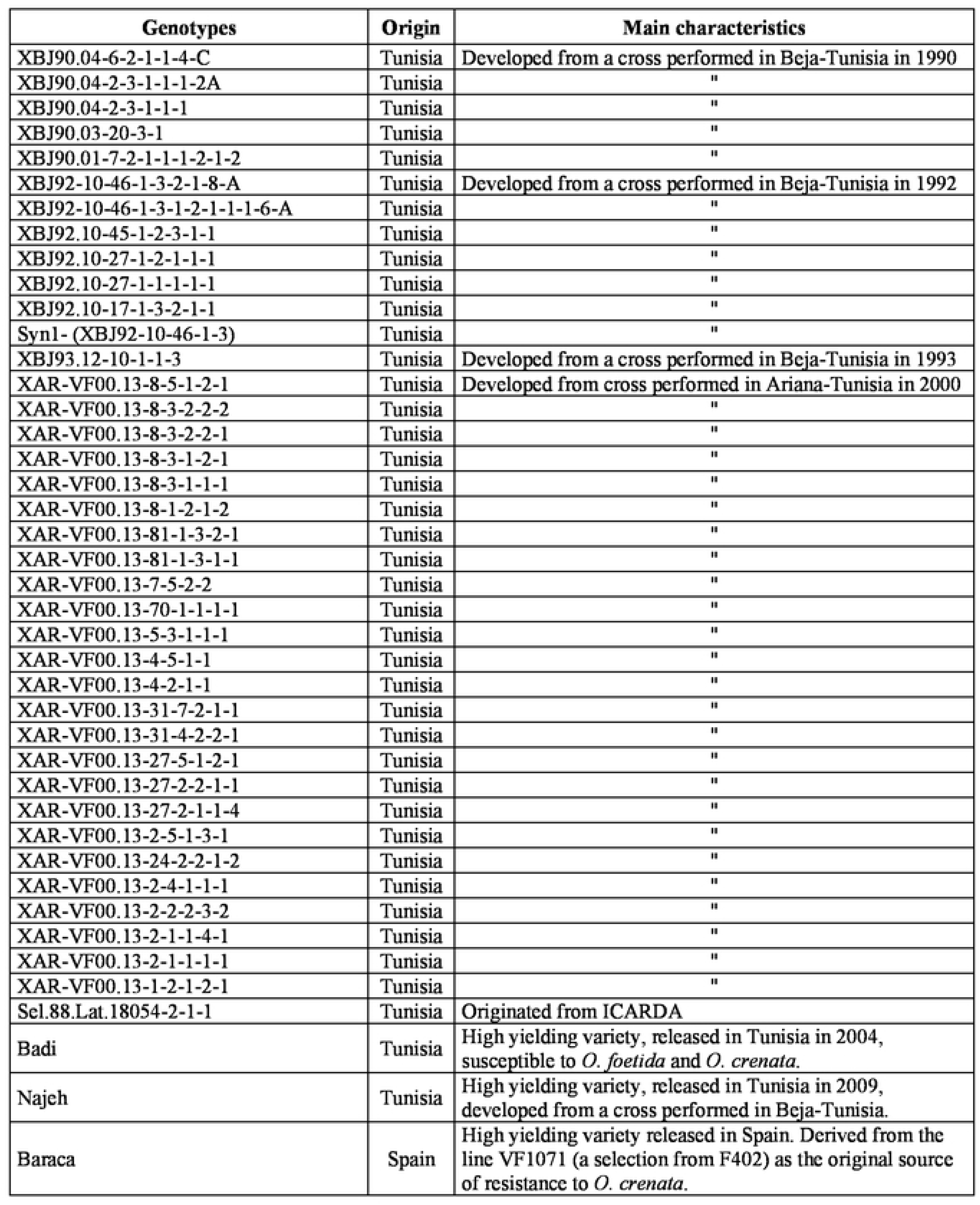
Origin and main characteristics of different studied genotypes

Out of the total tested collection, the two best resistant genotypes XAR-VF00.13-1-2-1-2-1 and XBJ90.04-2-3-1-1-1-2A were selected all with the three checks to conduct the second-year (2010/2011) evaluation and confirmation trial conducted under infested and non-infested fields. For both cropping seasons, trials were conducted in a randomized complete block design with three replications. Each genotype was planted, at a density of 24 seeds per m^2^, in four rows of 4 m length and 50 cm inter-rows spacing. The planting was performed the last week of November. No fertilizer’s supply or herbicide treatments were applied after plant emergence, only hand weeding was carried out. Monthly rainfall and average temperature distribution for the two cropping seasons collected from the iMETOS meteorological station (Pessl instruments) are presented in the Table 2.

**Table 2:** Climatic data (monthly minimum, maximum and average temperature (°C) and rain (mm) recorded in Beja research station during the two cropping seasons 2009/2010 and 2010/2011.

| Cropping season | Temp./Rain | Sep | Oct | Nov | Dec | Jan | Feb | Mar | Apr | May | Jun | Avg | Total |
| --- | --- | --- | --- | --- | --- | --- | --- | --- | --- | --- | --- | --- | --- |
| 2009-10 | Temp. min | 17.7 | 14.1 | 8.1 | 7.8 | 6.6 | 5.9 | 6.9 | 10.4 | 11.7 | 15.4 | 10.5 | - |
|  | Temp. max | 30.3 | 24.7 | 21.5 | 18.5 | 15.8 | 17.6 | 19.7 | 22.9 | 26.7 | 32.1 | 23.1 | - |
|  | Rain (mm) | 89.5 | 59.4 | 47.4 | 64.2 | 10.7 | 67.1 | 78 | 46.4 | 27.2 | 4.8 | - | 494.7 |
| 2010-11 | Temp. min | 17.3 | 13.6 | 10.7 | 6.2 | 5.6 | 5.1 | 6.4 | 9.5 | 12.3 | 15.2 | 10.2 | - |
|  | Temp. max | 30.7 | 26.7 | 20.8 | 17 | 15.9 | 15.5 | 18.3 | 23.5 | 26.4 | 31.5 | 22.6 | - |
|  | Rain (mm) | 42.7 | 82 | 56.8 | 61.6 | 63.2 | 138.4 | 58.3 | 38.6 | 43.4 | 7.4 | - | 592.4 |

### Measurements

The field response of the studied genotypes to *O. foetida* parasitism and their level of resistance was evaluated through different parameters that were measured at different host plant development stages.

During the first-year screening 2009/2010, at harvesting time data related to Parasitism Index (PI), number of emerged *Orobanche* shoots (EOS) per host plant and seed yield (SY) (g.m^−2^) were recorded. These data were recorded on the two central rows.

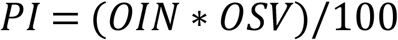

- OIN: *Orobanche* incidence or percentage of plants showing at least on *Orobanche* emerged shoot
- OSV: *Orobanche* severity (1-9 scale) or level of damage caused by *Orobanche* on the host plant development and seed production [24]

During the second-year evaluation 2010/2011, in addition to OIN, OSV, PI, EOS and SY mentioned earlier and recorded at harvesting time, other parameters were determined at pod-setting stage. from both infested and non-infested fields, five random plants from each plot were carefully dug-up and biomass and water content (WC) were recorded. Plants selected from infested field were dug-up with all *Orobanche* attachments that were later classified into emerged and non-emerged tubercles. Total emerged and non-emerged *Orobanche* number and dry weight per plant and number of days to *Orobanche* emergence (NDOE) were also recorded. Chlorophyll content index (CCI) was measured once a week between 10 am and 1 pm using a “*Hansatech*” CL-01 Chlorophyll Content Meter with a non-destructive method on leaves from the 11^th^ main stem node of five random host plants per plot. For every measurement almost the same part of the leaf was placed between two clips and the chlorophyll content index was determined in dual wavelength optical absorbance (620 and 940 nm).

The maximum quantum efficiency (F_v_/F_m_ ratio) was measured also once a week, before *Orobanche* emergence, in both infested and non-infested field between 10 am and 1 pm. from each plot, the measurements were performed on two random plants from the two central rows using a Plant Efficiency Analyzer (*Handy-PEA, Hansatech instruments Ltd, P02.002 v.*). For each plant, almost the same part/point of the leaflet situated on the 11^th^ main stem node was delimited by measure clip and was maintained in dark during 16 min by closing the clip shutter. Dark adaptation time was required to obtain a steady state value of the ratio of variable to maximum fluorescence. After 16 min, chlorophyll fluorescence transients were induced by a red light of 1500 μmol.m^−2^.s^−1^ intensity.

Plant sampling, biomass, water content (WC), Chlorophyll content index (CCI) and the maximum quantum efficiency (F_v_/F_m_ ratio) measurements were performed only during 2010/2011 cropping season.

### Statistical analysis

ANOVA was performed using the SPSS statistical program v.21 and differences among treatments for all measurements were compared at *P*=0.05 and using Duncan’s multiple-range test.

## Results

### Field evaluation and identification of potential resistance genotypes to O. foetida

In total, 39 faba bean genotypes all with resistant and susceptible checks were evaluated for their resistance to broomrape under high *O. foetida* infested field during the cropping season 2009/20010. Results showed high variability in the resistance to *O. foetida* between the genotypes. Significant differences between genotypes were observed for the number of emerged *Orobanche* shoots per plant (EOS), parasitism index (PI) and seed yield (SY) (fig 1). High negative correlation was observed between SY (r = 0.644, *p* ≤ 0.001) and both EOS (r = 0.753**) and PI (r = 0.770**). Almost 46% of the tested genotypes showed a resistance level higher than the resistant check Najeh. Both advanced lines XBJ90.04-2-3-1-1-1-2A and XAR-VF00.13-1-2-1-2-1 expressed a high resistance level to *O. foetida* with respective PI of 1.2 and 2.2. Only 0.9 and 1.4 EOS were observed for these two genotypes. Such resistance observed for these two genotypes was reflected by a high seed yield with 154.2 257g.m^−2^ and 257g.m^−2^ observed respectively for XBJ90.04-2-3-1-1-1-2A and XAR-VF00.13-1-2-1-2-1. They produced almost two (1.8) and three (2.9) times more than the resistant check Najeh.

**Figure 1:**
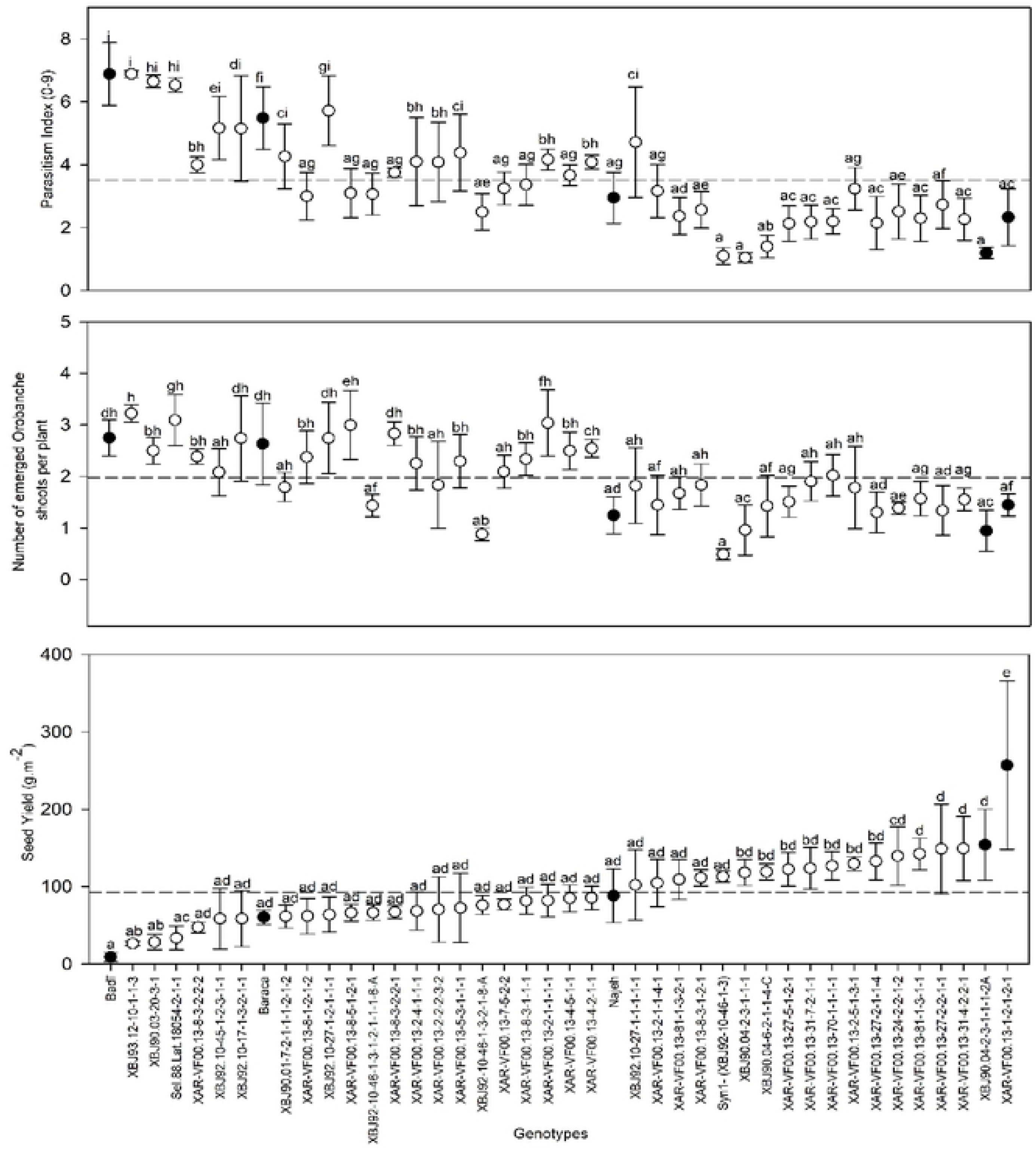
Parasitism Index (PI), Emerged Orobanche shoots per Plant (EOS) and Seed Yield (SY g.m^−2^) recorded for 39 genotypes under *O. foetida* infested field during the cropping season 2009/2010. Data are three replication means ± S E. Data with the same letter(s) are not significantly different at *P*=0.05 (Duncan test).

### Confirmation of the resistance under infested and non-infested field conditions

Among the 39 tested advanced lines during the cropping season 2009/2010, two lines XBJ90.04-2-3-1-1-1-2A and XAR-VF00.13-1-2-1-2-1 were selected for their high resistance *O. foetida.* These two lines, all with resistant and susceptible checks, were planted in 2010/2011 for further studies. The trial was conducted under infested and free *O. foetida* fields. ANOVA showed high differences (*P* ≤ 0.01) between the five studied genotypes for OIN, OSV, DOE, total *Orobanche* tubercles (TOT), and EOS. High OIN was observed for both cultivars Badi and Baraca with 81.7% and 85%, respectively (Table 3). Moderate incidence was observed for Najeh (65%) and the advanced line XAR-VF00.13-1-2-1-2-1 (60%). However, the advanced line XBJ90.04-2-3-1-1-1-2A showed the lowest *Orobanche* incidence (40%). Maximum infestation was observed for the susceptible genotype Badi with 5 tubercles per plant against only 1.2, 1.3 and 1.9 tubercles per plant observed on genotypes Najeh, XAR-VF00.13-1-2-1-2-1 and XBJ90.04-2-3-1-1-1-2A (Fig 2). A total *Orobanche* tubercles number of 4.1 (18.4% less than cv. Badi) was recorded for cv. Baraca which was reported to be resistant to *O. crenata* (Nadal et al., 2004). At crop maturity, the number of emerged *Orobanche* shoots (EOS) varied from 0.9 to 2.7 observed respectively for XBJ90.04-2-3-1-1-1-2A and Badi (Table 3). Only, 1.2, 1.4 and 2.6 shoots per plant were recorded respectively for Najeh, XAR-VF00.13-1-2-1-2-1 and cv. Baraca. *Orobanche* severity varied from 3 to 6.3 for XBJ90.04-2-3-1-1-1-2A and Badi, respectively. OSV of 4.3, 3 and 4.3 were recorded for the genotypes XAR-VF00.13-1-2-1-2-1, Najeh and Baraca, respectively (Table 3). Such infestation levels resulted in a significant negative parasitism impact on plant growth and seed production for the different tested genotypes. Differences between in the resistance to *O. foetida* were also confirmed by the number of days to *Orobanche* emergence which varied from 133 days for the susceptible check cv. Badi to 145 days observed for the advanced line XBJ90.04-2-3-1-1-1-2A (Table 3). Compared to Badi, a delay of 2.7, 4, 4.3 and 11.7 days was observed for DOE for the genotypes XAR-VF00.13-1-2-1-2-1, Baraca, Najeh and XBJ90.04-2-3-1-1-1-2A, respectively.

**Table 3:** *Orobanche* incidence(%) and *Orobanche* severity (1-9), Number of Days to Orobanche Emergence (DOE) and number of Emerged *Orobanche* Shoots per plant (EOS) recorded for different studied genotypes in high *O. foetida* infested field during the cropping season 2010/2011.

|  | *Orobanche* incidence (%) | *Orobanche* severity (1-9) | Number of days to *Orobanche* emergence | Emerged *Orobanche* shoot per plant at harvesting |
| --- | --- | --- | --- | --- |
| Badi | 100.0±0<sup>c</sup> | 6.3±1.2<sup>b</sup> | 133.0±2.6<sup>a</sup> | 2.7±0.6<sup>c</sup> |
| Baraca | 76.7±25.2<sup>bc</sup> | 4.3±1.2<sup>a</sup> | 139.7±6.1<sup>ab</sup> | 2.6±1.4<sup>b</sup> |
| Najeh | 50.0±30<sup>b</sup> | 3.0±0<sup>a</sup> | 141.7±5<sup>b</sup> | 1.2±0.6<sup>a</sup> |
| XAR-VF00.13-1-2-1-2-1 | 70.0±17.3<sup>bc</sup> | 4.3±1.2<sup>a</sup> | 140.3±3.1<sup>bc</sup> | 1.4±0.4<sup>a</sup> |
| XBJ90.04-2-3-1-1-1-2A | 13.3±5.8<sup>a</sup> | 3±0.0<sup>a</sup> | 145±2<sup>a</sup> | 0.9±0.7<sup>a</sup> |

Values followed by the same letter column are not significantly different at p=0.05 according to Duncan's multiple range mean comparison test

**Figure 2:**
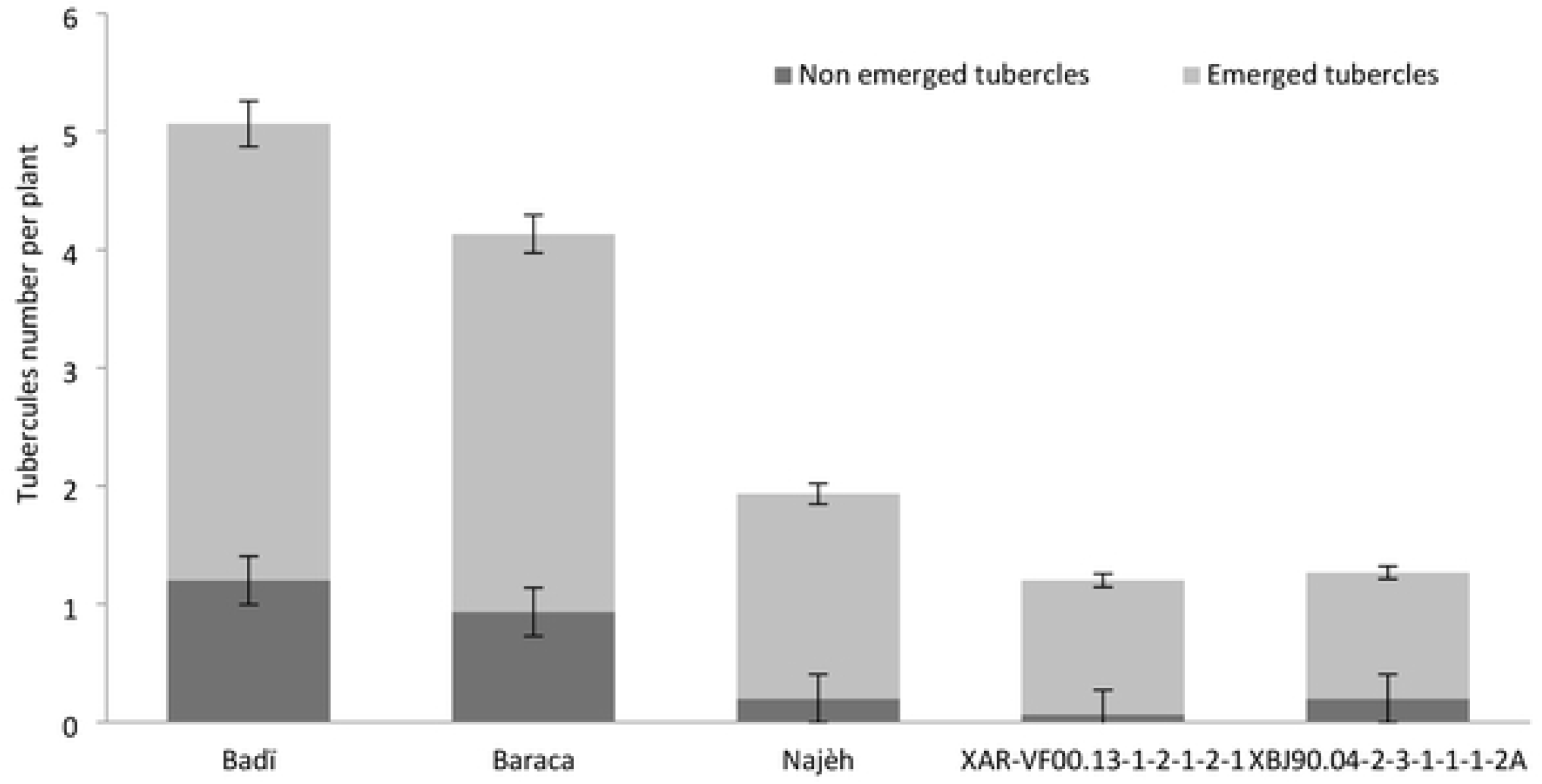
Total emerged and non-emerged *O. foetida* tubercles per plant recorded for different studied genotypes at pod setting stage. Data are three replication means± SE.

### Orobanche parasitism effect on biomass and seed yield

Compared to non-infested field, results showed that *O. foetida* has significantly affected the host plant biomass production (*P* ≤ 0.01) for all the studied genotypes (Table 4). A maximum decrease (68.5%) of biomass production was observed for the susceptible check Badi against only 15.6% recorded for Najeh. Respective decreases of 31%, 22.5% and 21.2% were recorded for the Baraca and both advanced lines XAR-VF00.13-1-2-1-2-1 and XBJ90.04-2-3-1-1-1-2A (Table 4). For all the studied genotypes, *Orobanche* parasitism has significantly affected biomass production but not the host plant water content (Table 4). Under infested conditions and contrary to biomass decreases, no significant reduction was observed for WC compared to non-infested plants. The susceptible check Badi showed the highest decrease (−8.5%).

**Table 4:**
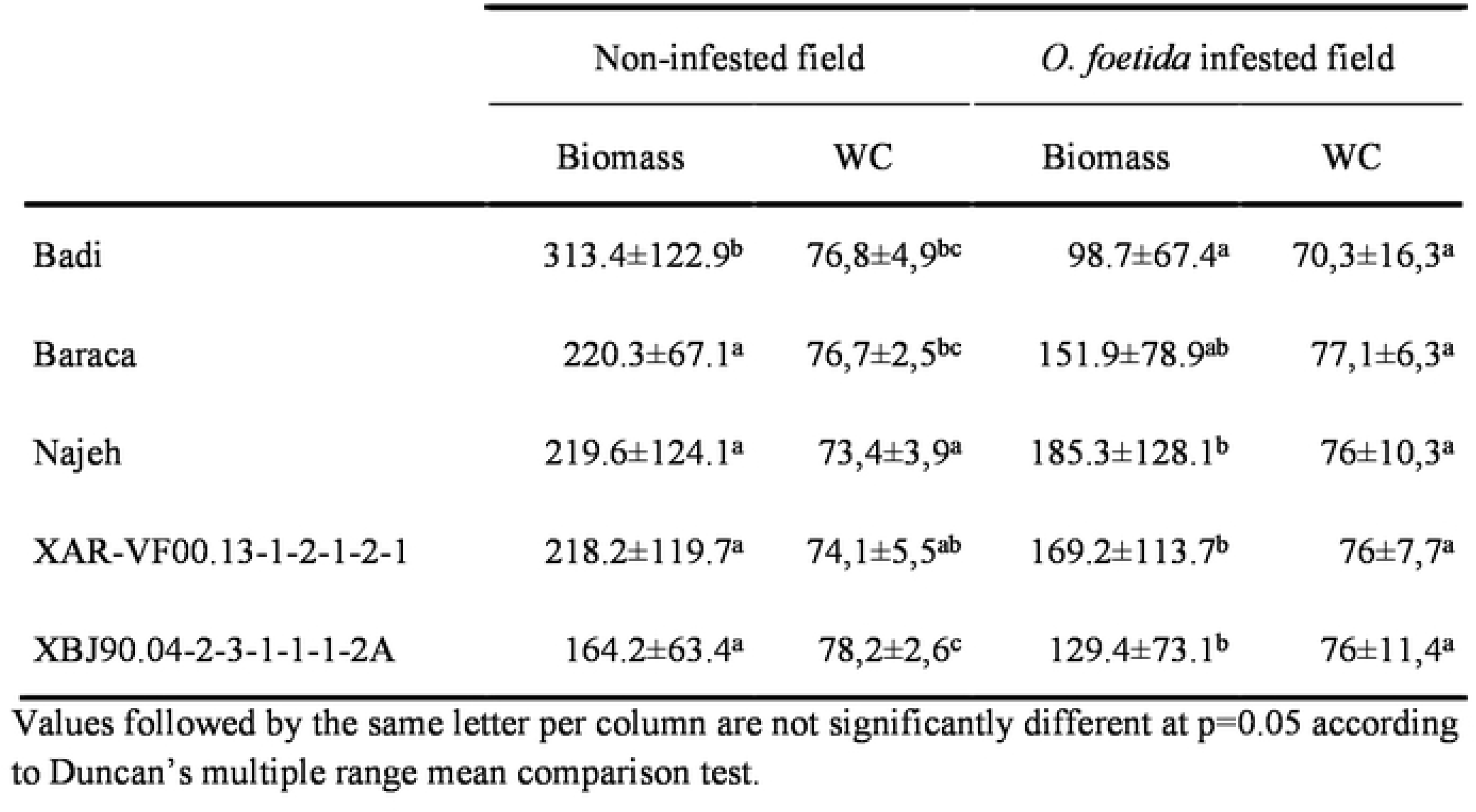
Biomass and water content (WC) recorded for different studied genotypes in both *O. foetida* infested and non-infested fields

|  | Non-infested field |  | *O. foetida* infested field |  |
| --- | --- | --- | --- | --- |
|  | Biomass | WC | Biomass | WC |
| Badi | 313.4±122.9<sup>b</sup> | 76,8±4,9<sup>bc</sup> | 98.7±67.4<sup>a</sup> | 70,3±16,3<sup>a</sup> |
| Baraca | 220.3±67.1<sup>a</sup> | 76,7±2,5<sup>bc</sup> | 151.9±78.9<sup>ab</sup> | 77,1±6,3<sup>a</sup> |
| Najeh | 219.6±124.1<sup>a</sup> | 73,4±3,9<sup>a</sup> | 185.3±128.1<sup>b</sup> | 76±10,3<sup>a</sup> |
| XAR-VF00.13-1-2-1-2-1 | 218.2±119.7<sup>a</sup> | 74,1±5,5<sup>ab</sup> | 169.2±113.7<sup>b</sup> | 76±7,7<sup>a</sup> |
| XBJ90.04-2-3-1-1-1-2A | 164.2±63.4<sup>a</sup> | 78,2±2,6<sup>c</sup> | 129.4±73.1<sup>b</sup> | 76±11,4<sup>a</sup> |

Values followed by the same letter per column are not significantly different at p=0.05 according to Duncan's multiple range mean comparison test.

*Orobanche* parasitism effect on host plant biomass was reflected by seed yield losses for all the studied genotypes. Compared to non-infested plants, seed yield decreases varied from a minimum of 3.9% recorded for XBJ90.04-2-3-1-1-1-2A to a maximum of 93.9% observed for the susceptible check Badi. Respective decreases of 77.4%, 39.5% and 28.8% were recorded for the genotypes Baraca, Najeh and XAR-VF00.13-1-2-1-2-1 (Fig 4). Among all the tested genotypes, XAR-VF00.13-1-2-1-2-1 was the most productive under *Orobanche* infested conditions with 228.4 g.m^−2^ representing 3 and 4 times the seed yield recorded for the resistant checks Najeh (78.3 g.m^−2^) and Baraca (53.8 g.m^−2^). Seed production for cv. Baraca which is reported to be resistant to *O. crenata* varied from 237.8 g.m^−2^ to 53.8 g.m^−2^ (77.4% less) under free and *Orobanche* infested fields, respectively.

**Figure 3:**
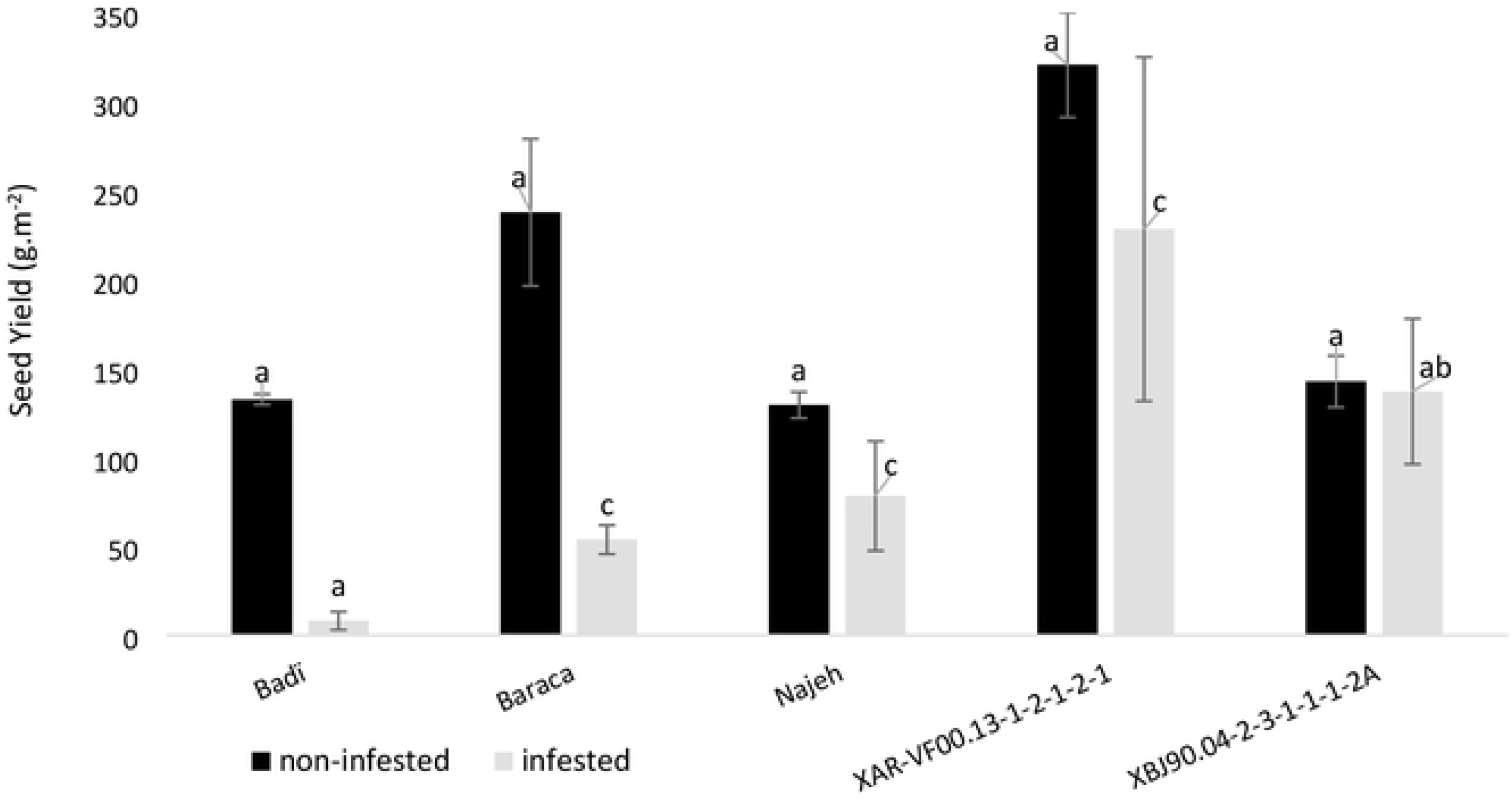
Seed yield (g.m^−2^) recorded for different studied genotypes in both *O. foetida* infested and non-infested fields during the cropping season 20I0/2011. Data are three replication means ± SE. For each treatment (infested and non-infested) data with the same letter(s) are not significantly different at *P*=0.05 (Duncan test).

**Figure 4:**
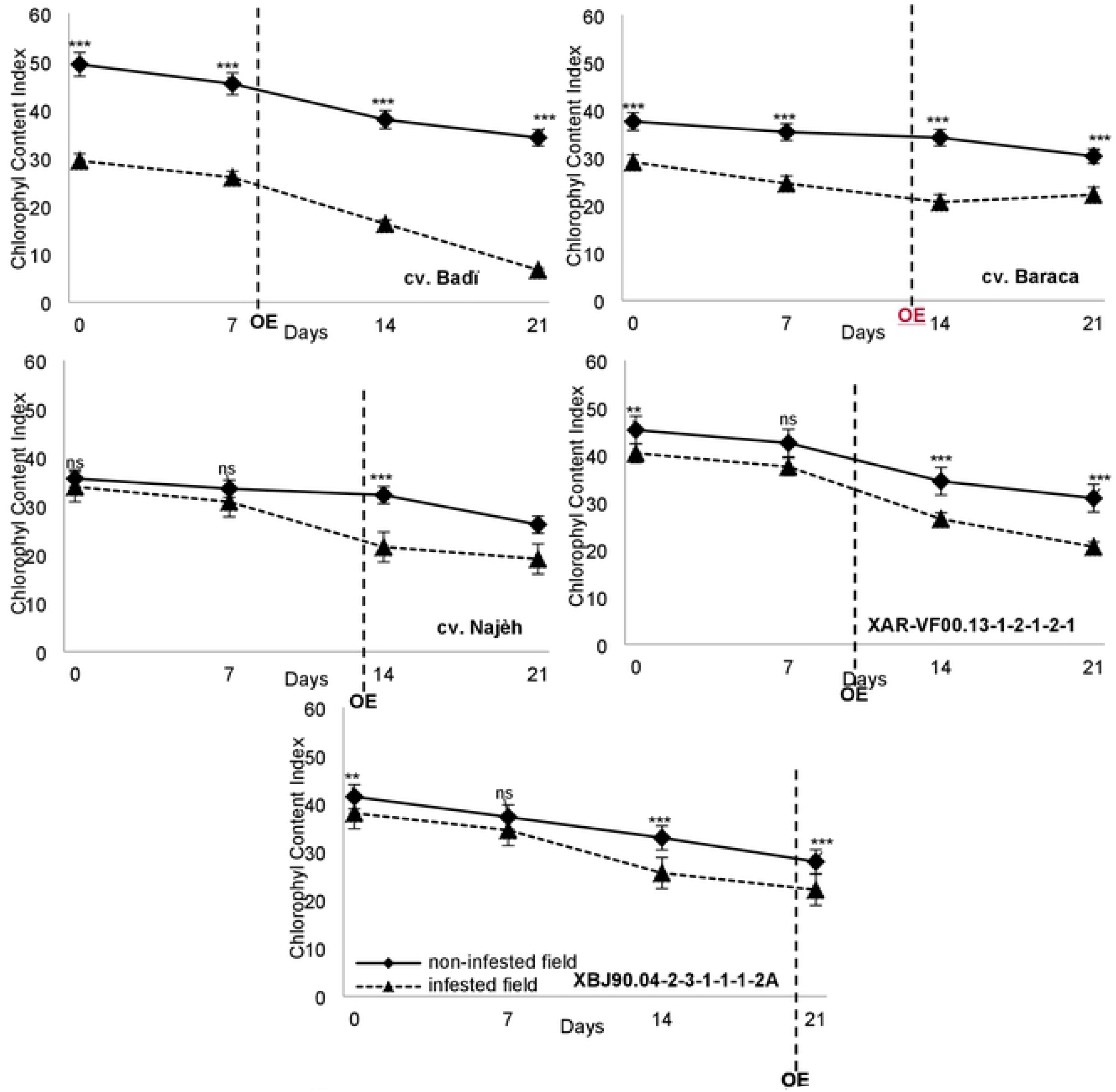
Chlorophyll content index (CCI) recorded for the different studied genotypes in both *O. foetida* infested and non-infested fields during the cropping season 2010/2011. OE: *Orobanche* Emergence. Data are fifteen replications means ± SE. Data with the same letter(s) per are not significantly different at P=0.05 (Duncan test).

### Chlorophyll Content Index and chlorophyll fluorescence

Results showed that compared to non-infested plants (free *Orobanche* field), *Orobanche* parasitism highly affected (*P* ≤ 0.01) the host plant chlorophyll content index (CCI) and F_v_/F_m_ ratio for all the studied genotypes (Fig 4 and 5).

**Figure 5:**
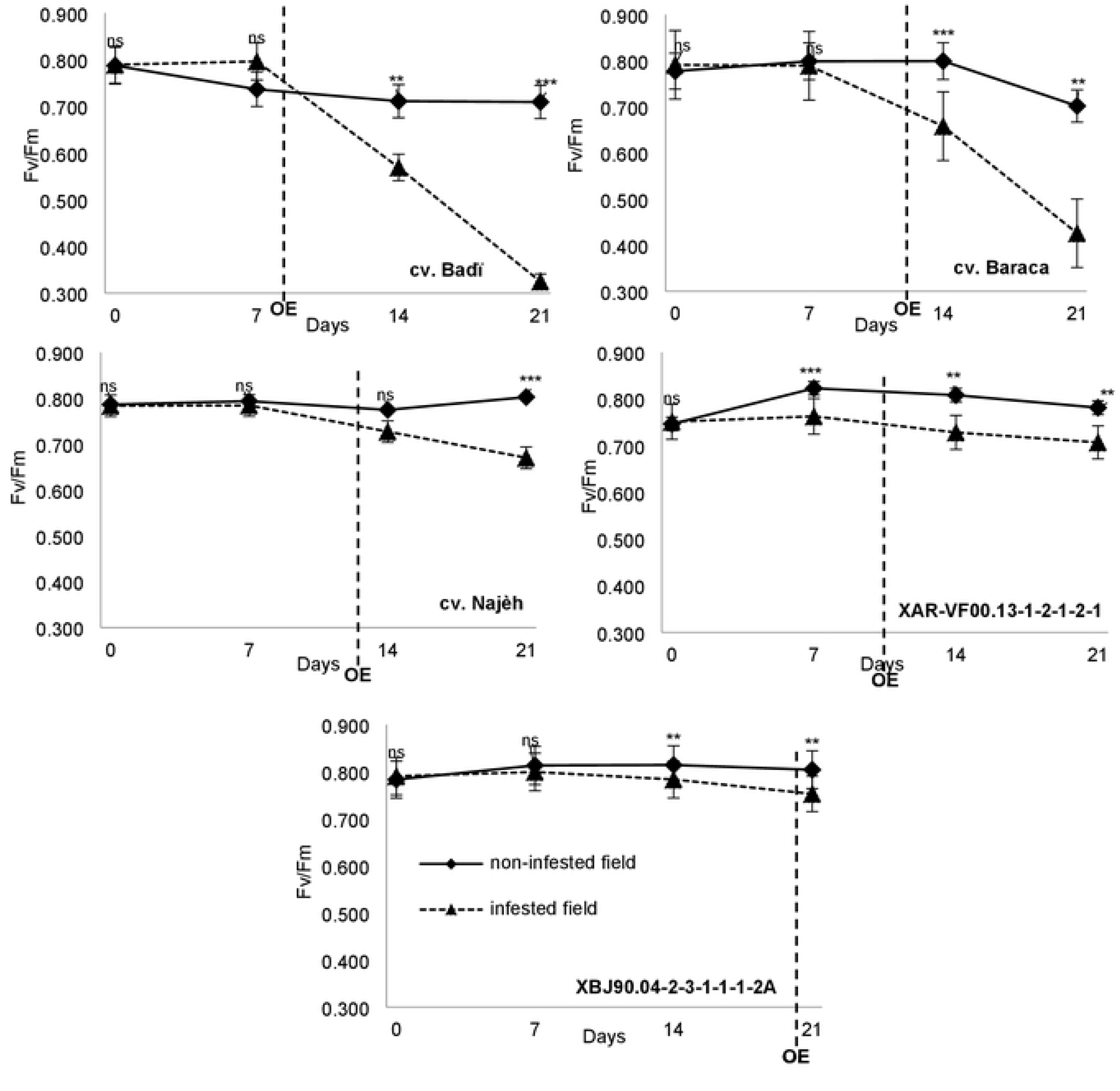
F_v_/F_m_ ratio recorded for the different studied genotypes in both *O. foetida* infested and non-infested fields during the cropping season 2010/201 I. OE: *Orobanche* Emergence. Data are 6 replications means ± SE. ns: non significantly different at P≥0.05; ***: significantly different at P≤0.01; **: significantly different at *P*≤0.05.

Under infested conditions, CCI decreases varied from 23.6% for Baraca to 77.2% recorded for the susceptible check Badi between the first and last scoring dates. For the same genotypes respective decreases of 19.4% and 30.8% were observed under free *Orobanche* field. Under infested conditions, decreases of 43.8%, 49% and 41.9% were recorded respectively for the other genotypes Najeh, XAR-VF00.13-1-2-1-2-1 and XBJ90.04-2-3-1-1-1-2A, against only 26.6%, 31.8% and 32.5% recorded in non-infested field. Clear differences were observed for the CCI between infested and non-infested plants. Such differences were highly significant and more pronounced for the susceptible check Badi and Baraca. Differences in the variation between 1^st^ and last score under infested and non-infested conditions varied from minimum of 4.2% for Baraca and maximum of 46.4% observed for the susceptible check Badi. A difference of 17.2% was recorded for both Najeh and XAR-VF00.13-1-2-1-2-1 and 9.3% observed for XBJ90.04-2-3-1-1-1-2. Except Najeh, all the other genotypes showed high significant difference between infested and non-infested plants before *Orobanche* emergence.

Results, also, showed that, at the end of the experiment and for all tested genotypes, the maximum quantum efficiency (F_v_/F_m_ ratio) was significantly affected by *Orobanche* parasitism (Fig 5). Under infested field conditions, F_v_/F_m_ decreased by 58.8% (0.789 to 0.325) for the susceptible check Badi against only 9.9% (0.787 to 0.709) observed for non-infested plants. Decreases of 46.2%, 14.5%, 5.9% and 4.7% were recorded, respectively, for the genotypes Baraca, Najeh, XAR-VF00.13-1-2-1-2-1 and XBJ90.04-2-3-1-1-1-2A. No significant variation of F_v_/F_m_ was observed for all the studied genotypes under free *Orobanche* conditions. For the three checks Badi, Baraca and Najeh, and before *Orobanche* emergence, no significant differences were observed in F_v_/F_m_ ratio between infested and non-infested plant (fig 5). For both selected genotypes XAR-VF00.13-1-2-1-2-1 and XBJ90.04-2-3-1-1-1-2A significant differences were recorded between infested and non-infested plants before *Orobanche* emergence. High decreases of F_v_/F_m_ ratio were observed before *Orobanche* emergence for both genotypes Badi and Baraca against only slight decreases recorded for Najeh and both selected advanced lines XAR-VF00.13-1-2-1-2-1 and XBJ90.04-2-3-1-1-1-2A. At the end of the experiment comparison of F_v_/F_m_ ratio between non-infested and infested plants showed differences of 6.4% (0.804 vs 0.753) and 9.5% (0.781 vs 0.707) for both genotypes XBJ90.04-2-3-1-1-1-2A and XAR-VF00.13-1-2-1-2-1 against 54.2% (0.709 vs 0.325), 39.3% (0.702 vs 0.426) and 16.5% (0.802 vs 0.670) recorded for the susceptible and resistant checks Badi, Baraca and Najeh, respectively.

## Discussion

Results from the first-year screening showed high variability for the resistance to *O. foetida* in the tested collection. Two advanced lines, XBJ90.04-2-3-1-1-1-2A and XAR-VF00.13-1-2-1-2-1 were identified and selected for their high resistance level and high yield under heavy *O. foetida* infested conditions. Confirmation trials conducted under infested and non-infested conditions using both genotypes and susceptible and resistant checks during the two cropping seasons 2009/2010 and 2010/2011 revealed high significant variation (*P* ≤ 0.01) between the studied genotypes in term of resistance to *O. foetida*. In general, resistance to broomrapes is, not only the capability of the host to limit the parasite development and the damages that causes, but also the capacity of the host to grow and produce grains under such parasitism attack. Significant differences were recorded for DOE, TOT, EOS and *Orobanche* incidence and severity. Under *Orobanche* infested conditions and compared to the susceptible check Badi, the genotypes Baraca, Najeh, XAR-VF00.13-1-2-1-2-1 and XBJ90.04-2-3-1-1-1-2A showed a low infestation level. Najeh and both advanced lines XAR-VF00.13-1-2-1-2-1 and XBJ90.04-2-3-1-1-1-2A showed the highest resistance to *O. foetida* while Baraca, previously reported to be resistant to *O. crenata* [23], expressed a moderate resistance to *O. foetida*. Furthermore, data showed that for all five genotypes, biomass and seed production were negatively affected by the parasite. Compared to non-infested plants, early wilting symptoms were observed for parasitized plants resulting in a shortening of the reproductive phase and affecting the plant biomass and grain yield. For Badi, *Orobanche* has severely restrained plant growth, affected the flowering and pod setting, and resulted in almost complete damage and yield losses. A moderate effect of the parasite on plant development and seed production was observed for other tested genotypes Baraca, Najeh, XAR-VF00.13-1-2-1-2-1 and XBJ90.04-2-3-1-1-1-2A. Results also showed that despite the biomass decreases recorded for different studied genotypes, no significant effect of *Orobanche* parasitism was observed on the host water content (WC) which varied from 70.3% to 78.2% under both free and infested conditions. This could be explained by the fact that due to the parasitic burden and resources sinking the host plant limited its biomass and dry matter production and allocation in order to keep its physiological functioning through a normal and optimum water content. *Orobanche* parasitism effects on host plant growth and biomass production and allocation are directly related to the infestation level. Ennami et al., [25] reported a high negative correlation between faba bean and lentil plant growth and biomass production and the number and size of *Orobanche* shoots/tubercles and thus the nutrients sinking level. Other previous studies reported that the detrimental effect of both *O. foetida* and *O. crenata* on faba bean grain yield can reach up to 90% - 100% depending on infection severity and the broomrape-crop association [26, 27]. Furthermore, Cameron et al. [28] reported that the parasitic plant *Rhinanthus minor* significantly reduced biomass production in *Phleum bertolinii* and demonstrated that such decrease was reflected by changes in photosynthetic activities and significant reductions in the quantum efficiency of PS II and chlorophyll concentration. In fact, the number of leaves of host plants as well as leaf greenness is very important in plant eco-physiological studies because it provides information about physiological responses of plants under stress conditions [29, 30]. In our study, results showed that compared to the non-infested plants, CCI was significantly affected by *O. foetida* for the five studied genotypes. Significant differences in CCI were observed, between infested and non-infested plants, before *Orobanche* emergence which can make this parameter very useful for early detection of the underground *Orobanche* infestation. In addition, decreases in CCI for infested plants could be explained by the parasite nutritional requirements that limits the normal growth and functioning of the host plant. Similar results were reported for tomato/*P. ramosa* pathotype [31] and *Mikania micrantha*/*Cuscuta campestris* [20]. The latter study showed that despite the CCI decrease observed on the *M. micrantha* leaves, there was no significant effects of *C. campestris* parasitism on chlorophyll a:b ratio.

Our results demonstrated also that CCI decreases was associated with photosynthetic characteristics variation in the host plant leaves. *O. foetida* affected the photosynthetic system through significant decreases of the leaves CCI and the maximum quantum efficiency (F_v_/F_m_ ratio) which was increasingly pronounced over time, especially for the susceptible check Badi.

For the different studied genotypes, *O. foetida* parasitism effect on faba bean plants resulted in a significant (*P* ≤ 0.01) decrease of the F_v_/F_m_ ratio as compared to non-infested plants. High significant difference was observed in F_v_/F_m_ ratio between infested and non-infested plant before *Orobanche* emergence for both advanced lines XAR-VF00.13-1-2-1-2-1 and XBJ90.04-2-3-1-1-1-2A. Despite *Orobanche* parasitism effect, these two genotypes were able to maintain a good functioning of their PSII as normal as for the free-orobanche plants even after *Orobanche* emergence. Contrary, this was not the case for the three susceptible and resistant checks Badi, Baraca and Najeh as no significant differences in F_v_/F_m_ ratio between infested and non-infested plant before *Orobanche* emergence. These genotypes maintained normal functioning but under an increasing parasitism pressure, important decreases in F_v_/F_m_ ratio were recorded at *Orobanche* emergence for three genotypes, especially, Badi and Baraca. These results all with the analyses of F_v_/F_m_ ratio recorded for all five genotypes in both free and infested *O. foetida* fields, indicated that F_v_/F_m_ ratio could be used not only for the quantification of stress caused by *Orobanche* parasitism and early detection of the underground infestation but also the screening and identification of high resistant genotypes.

Similar results were reported by Mauromicale et al. [31] who showed that F_v_/F_m_ ratio, which is proportional to the PS II quantum yield and well correlated with the photosynthesis quantum yield [32], was significantly reduced by *P. ramosa* attack on tomato. In the same study, the authors demonstrated that the F_v_/F_m_ reduction is mainly induced by an effect on the variable fluorescence (F_v_) resulted on damage in PS II electron transport. In addition, Jeschke and Hilpert [33] showed that *Cuscuta reflexa* induced a sink-dependent stimulation of net photosynthesis on *Ricinus communis*. Shen et al. [20] showed that *C. campestris* infection decreases host stomatal conductance, transpiration, chlorophyll content, and soluble protein concentration on *M. micrantha*, which may directly and indirectly reduce the photosynthesis rate and affect the host plant growth. These results are contrasting with other studies [34, 35] who reported that broomrape affects host biomass and yield and related traits with only minor disturbance to leaves tissue but no perceptible effects on photosynthetic rate. More recently, Ennami et al. [25] showed that effective quantum yield of open photosystem II, (Fm’-F)/Fm’, was significantly reduced by *O. crenata* attack on susceptible faba bean and lentil genotypes.

## Conclusions

Results showed that *O. foetida* can affect faba bean host plants in/through different ways and at a big range of scales, from the root to the leaves through the whole plant. Compared to non-infested plant, high significant difference was recorded between different studied genotypes in response to *O. foetida* parasitism. The genotypes Najeh, XAR-VF00.13-1-2-1-2-1 and XBJ90.04-2-3-1-1-1-2A which expressed the highest resistance levels to *O. foetida* showed a moderate decrease of biomass and seed production. A significant variation in CCI and F_v_/F_m_ ratio was observed from individual plants between the tested genotypes and between free and infested plants for the same genotype. The significant positive correlation observed between F_v_/F_m_ ratio and high resistance level to *O. foetida* may suggest that this physiological parameter could be potentially used as a practical screening tool integrating physiological trait for plant selection and early detection of the root parasitic weeds infestation.

## Acknowledgment and funding support

This work was supported by the Tunisian Ministry of Agriculture, Hydraulic Resources and Fishery and the Ministry of Higher Education and Scientific Research. The authors are thankful to all the field crop laboratory’s technical staff for their assistance; Leila Dakhli, Fadhel Sellami, Hadhemi Abidli, Besma Soltani and Olfa Mlayeh.

## Author’s contribution

M.A., Z.A., I.T., M.K., designed the research. M.A., Z.A., M.K., performed the experiments. M.A., Z.A., I.T., R.M., M.E.G., M.K., contributed materials/analysis tools. M.A., Z.A., wrote the paper. I.T., M.E.G., R.M., M.K., revised the paper. All authors approved the final manuscript.

